# 3D genome evolution and reorganization in the *Drosophila melanogaster* species group

**DOI:** 10.1101/2020.04.09.033753

**Authors:** Nicole S. Torosin, Aparna Anand, Tirupathi Rao Golla, Weihuan Cao, Christopher E. Ellison

## Abstract

Topologically associating domains, or TADs, are functional units that organize chromosomes into 3D structures of interacting chromatin. TADs play an important role in regulating gene expression by constraining enhancer-promoter contacts; there is evidence that deletion of TAD boundaries leads to aberrant expression of neighboring genes. While the mechanisms of TAD formation have been well-studied, current knowledge on the extent of TAD conservation across species is inconclusive. Due to the integral role TADs play in gene regulation, their structure and organization is expected to be conserved during evolution. However, more recent research suggests that TAD structures diverge relatively rapidly. We use Hi-C chromosome conformation capture to measure evolutionary conservation of whole TADs and TAD boundary elements between *D. melanogaster* and *D. triauraria*, two early-branching species from the *melanogaster* species group which diverged ~15 million years ago. We find that 75% of TAD boundaries are orthologous while only 25% of TAD domains are conserved and these are enriched for Polycomb-repressed chromatin. Our results show that TADs have been reorganized since the common ancestor of *D. melanogaster* and *D. triauraria*, yet the sequence elements that specify TAD boundaries remain highly conserved. We propose that evolutionary divergence in 3D genome organization results from shuffling of conserved boundary elements across chromosomes, breaking old TADs and creating new TAD architectures. This result supports the existence of distinct TAD subtypes: some may be evolutionarily flexible while others remain highly conserved due to their importance in restricting gene-regulatory element interactions.

## 1 Introduction

The recent development of Hi-C sequencing techniques has allowed inference of three-dimensional chromosome conformation through identification of inter- and intra-chromosomal interactions at high-resolution, across the entire genome. Visualization of gene contacts and contact frequencies led to the discovery of organizational features called topologically associating domains, or TADs, which bring genes in close proximity with their regulatory elements [41]. TADs are regions of highly interacting chromatin that contain genes with similar expression patterns and epigenetic states [12, 61]. Domains are demarcated by boundaries which are regions of decompacted chromatin bound by insulator proteins [63]. In vertebrates, the CCCTC-binding factor (CTCF), along with the structural maintenance of chromosomes (SMC) cohesin complex, play a major role in specifying TAD boundaries [56, 12, 68, 52], whereas in Drosophila, CTCF and SMC binding show little enrichment at TAD boundaries. Instead, other insulator proteins, including BEAF-32, Chromator, CP190, and M1BP are more frequently found at TAD boundaries [61, 55, 31, 30, 67] and depletion of M1BP has been shown to disrupt 3D genome organization in the Drosophila Kc167 cell line [55].

Most research thus far investigating 3D genome structure has operated under the prevailing theory that TADs regulate gene expression by limiting potential gene-enhancer interactions to those within a given domain. This theory is supported by a variety of studies. For example, genome-wide enhancer-promoter contacts in mouse neurons occur mainly within TADs [6] and reporter gene-enhancer interactions have been shown to be correlated with TAD structure [65]. Furthermore, disruption of TAD boundaries has been associated with aberrant enhancer/promoter contacts, gene misregulation, developmental abnormalities and cancer [43, 22, 21, 29, 44, 70].

The functional role of TADs with respect to the regulation of gene expression has important implications for 3D genome evolution. If TAD structure is critical for proper gene regulation, then the evolution of 3D genome organization should be highly constrained and related species should show strong conservation of TAD structures. Consistent with this prediction, a variety of studies in vertebrates have reported strong conservation of 3D genome organization using comparative Hi-C approaches [12, 68, 35, 38]. A recent study in Drosophila reported that 3D genome architecture is conserved over 40 million years of evolution in spite of extensive chromosomal rearrangements [57]. These studies support a model where chromosomal rearrangements that preserve TADs (i.e. their breakpoints are located within TAD boundaries) are much more likely to be retained over evolutionary time compared to rearrangements that disrupt TADs (Figure 1, Model 1). Under this model, TADs are shuffled as whole units over evolutionary time due to selection to maintain the 3D interaction properties of the genes and regulatory sequences within them.

**Figure 1:**
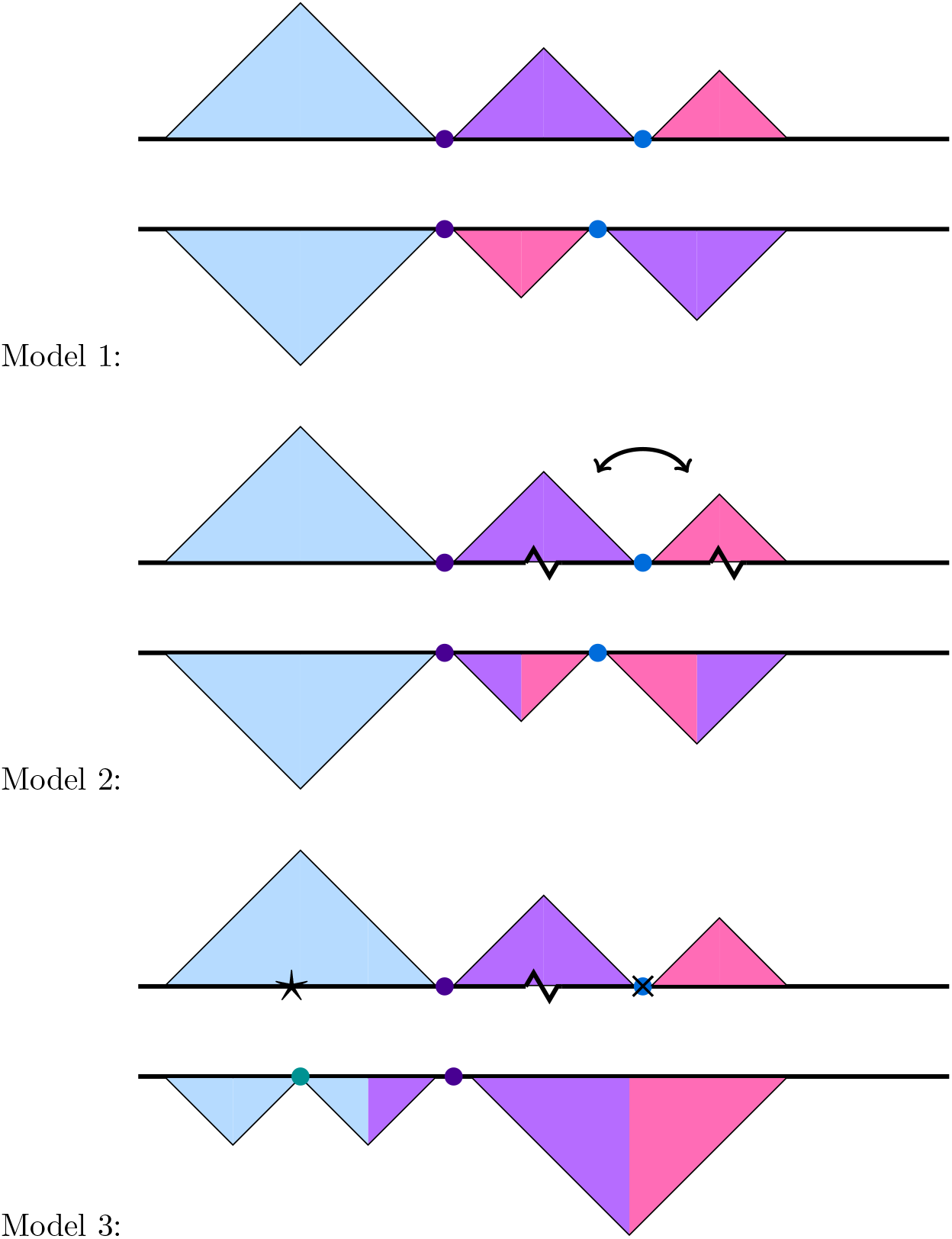
Models depicting possible TAD and boundary rearrangement scenarios. Each model shows TAD contact domains (triangles) and boundaries (circles) in two hypothetical species. Model 1: chromosomal rearrangements occur at TAD boundaries, resulting in domains shuffling as intact units, represented by the purple and lilac TADs. Model 2: TAD boundaries remain conserved but TADs themselves are disrupted. In this example, an inversion whose breakpoints (jagged lines) occur within two separate TADs results in TAD reorganization, seen here as the two chimeric purple/lilac TADs. Model 3: both TADs and their boundaries evolve rapidly. The gain and loss of boundary elements (black star and ‘X’, respectively), leads to further TAD reorganization, beyond that observed in Model 2.

However, more recent research suggests that TADs may be frequently reorganized over evolutionary time. Notably, one recent study found that only 43% of TADs are shared between humans and chimpanzees [19]. Furthermore, there are a number of studies showing that gene expression profiles remain unperturbed upon TAD reorganization. For example, the extensive changes in chromosome topology caused by the rearrangements found on a Drosophila balancer chromosome are not associated with differences in gene expression [24]. Also in Drosophila, studies involving deletion mutations [39] and experimentally-induced inversions [47] found that these mutations, which should disrupt TAD organization, had little effect on gene expression. Similar observations have been made in mice: fusion of TADs does not have major effects on gene expression [10]. These studies suggest that TADs may diverge relatively rapidly over evolutionary time, with little effect on gene expression (see Models 2 and 3, Figure 1).

There are several possibilities that might explain these apparently contradictory results. Some studies [12] only assess the conservation of TAD boundaries, rather than TADs themselves. It is possible that boundary elements evolve at a different rate than TADs. It is also possible that there are distinct functional subtypes of TADs, with some being more tolerant of reorganization than others. Consistent with such a possibility, a recent study identified a subset of ancient, highly-conserved TADs in both vertebrates and flies that are enriched for conserved noncoding elements and developmental genes [26].

Here, we have compared 3D genome organization between *Drosophila melanogaster* and *Drosophila triauraria*, which diverged ~15 million years ago [50]. We chose this species pair because they have accumulated extensive chromosomal rearrangements since their divergence, yet maintain blocks of synteny that are at least twice as large as TADs (~120 Kb on average, see Results, Drosophila TADs: 23 - 63 Kb average length [55]). We have improved a previously published *D. triauraria* genome assembly [48] by performing additional nanopore sequencing and Hi-C scaffolding which yielded chromosome-length scaffolds. We then used two biological replicates of Hi-C sequencing data to identify high-confidence TADs and TAD boundaries in each species. We separately assessed the evolutionary conservation of boundary elements and complete TADs and found that TAD boundaries are significantly more conserved between these two species than the TADs themselves. Overall, we find that only 25% of TADs are orthologous and these conserved TADs are enriched for Polycomb-repressed chromatin, similar to what has been observed by Harmston et al. [26]. Our results show that TADs have been reorganized since the common ancestor of *D. melanogaster* and *D. triauraria*, yet the sequence elements that specify TAD boundaries remain highly conserved. We propose that evolutionary divergence in 3D genome organization results from shuffling of conserved boundary elements across chromosomes, breaking old TADs and creating new TAD architectures. Our results also support the existence of functionally distinct TADs subtypes: many TADs may be evolutionarily flexible and able to be reorganized without perturbing gene expression, whereas there may also be a distinct set of TADs subtypes that remain highly conserved due to their importance in restricting long-distance gene-regulatory element interactions.

## 2 Results

### 2.1 *D. triauraria* genome assembly

A recently published genome assembly for *D. triauraria* was made using relatively low-coverage (~18.8×) nanopore sequencing data [48]. In order to create an improved assembly, we performed additional long-read nanopore sequencing of genomic DNA extracted from ~30 adult females from *D. triauraria* strain 14028-0651.00 (National Drosophila Species Stock Center at Cornell). We used three r9.4 flow cells to generate a total of 633,844 reads (10,287 bp mean length) and 6.5 Gb of sequencing data. We combined our data with the previously published nanopore data from *D. triauraria* [48] for a final dataset of 1,043,600 reads (total size: 10.5 Gb, coverage: 52×). We basecalled the raw signal data with *Albacore* (available from Oxford Nanopore Technologies) and assembled the basecalled reads with *Canu* [33], which produced an assembly with contig N50 of 1.3 Mb (1098 total contigs, 269 Mb total size). We then polished the assembly by using *Nanopolish* [42] with raw nanopore signal data and *Pilon* [69] with Illumina data, which corrected a total of 1,185,510 assembly errors. Next, we used *Purge Haplotigs* [58] to identify allelic contigs, where highly heterozygous haplotypes were assembled as separate contigs rather than collapsed. After removing secondary haplotigs and bacterial contigs, our final contig assembly consisted of a total of 294 contigs which sum to ~200 Mb and have an N50 of 1.7 Mb, which is 2.4× larger than the previously published study (N50 = 720 Kb) [48] (Table S1).

We next performed Hi-C scaffolding using the polished nanopore contigs and the software packages *Juicer* [15] and *3D-DNA* [13] (Figure 2a,b). The scaffolding process produced chromosome-length scaffolds, reflected by the dramatic increase in N50 from 1.7 Mb to 34.5 Mb. In order to assign the *D. triauraria* scaffolds to Muller elements, we performed a translated BLAST search of our scaffolds using *D. melanogaster* peptides as queries and keeping only the best hit for each query sequence. We found that each scaffold was highly enriched for *D. melanogaster* peptides from a specific Muller element (Figure S1) and we successfully identified scaffolds corresponding to Muller elements A-F for downstream analysis. In order to predict complete gene models for our assembly, we generated RNA-seq data from *D. triauraria* ovaries, testes, and embryos. We combined the RNA-seq data with *ab initio* gene predictions from *Augustus* [64] and *SNAP* [34] and homology-based predictions from *D. melanogaster* peptides using *MAKER* [8].

**Figure 2:**
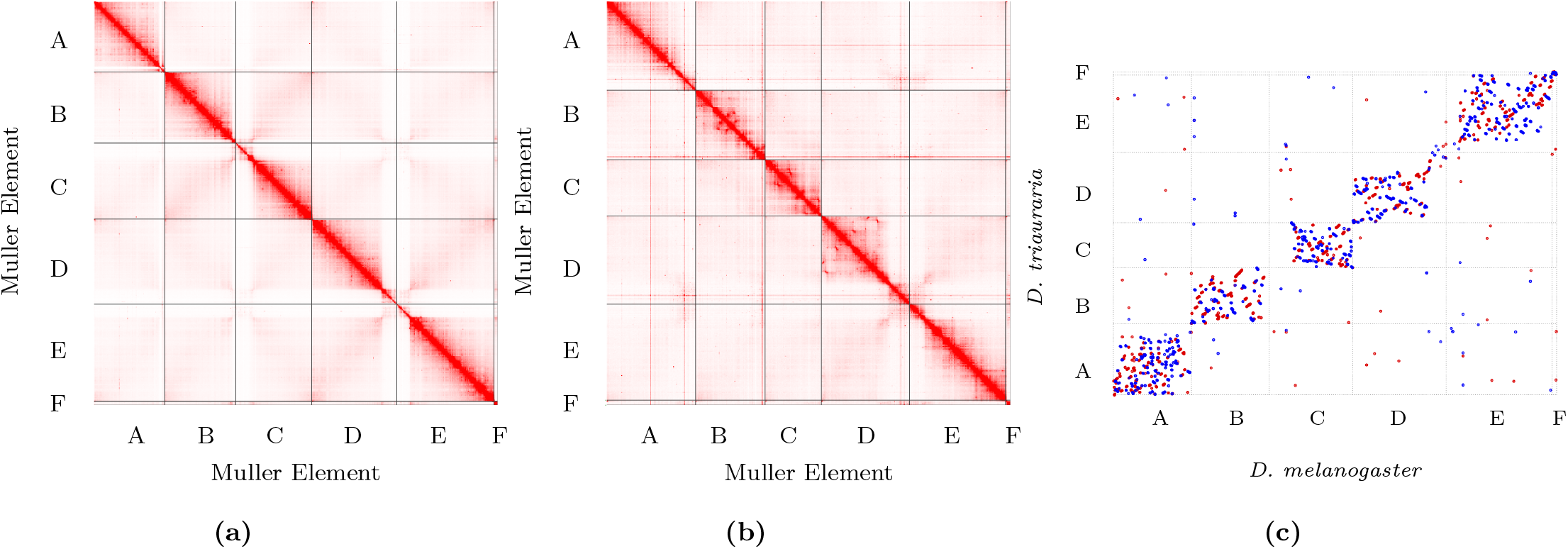
Hi-C contact maps and genome comparison. a) *D. melanogaster* contact map; b) *D. triauraria* contact map. Contact maps show frequencies of pairwise 3D contacts, inferred from Hi-C data. Darker colors represent higher contact frequencies. Contact frequencies were visualized using *Juicebox* [15]. c) *MUMmer* dotplot depicting chromosomal rearrangements between *D. melanogaster* and *D. triauraria*. The promer utility from *MUMmer* [36] was used to visualize synteny between *D. melanogaster* and *D. triauraria*. Each dot corresponds to a one-to-one alignment between the two genomes. Red dots represent +/+ strand alignments and blue dots represent +/− strand alignments.

*D. triauraria* was previously sequenced unintentionally because it was mislabeled as *D. kikkawai* [48], which means that the published *D. triauraria* nanopore data are from an unknown strain. The strain we sequenced is from the same stock center, making it likely that the contaminant and our strain, are in fact the same strains. To test this, we aligned the Illumina data from Miller et. al. [48], in conjunction with our uninformative Hi-C reads, to our nanopore assembly and called SNPs using *FreeBayes* [23]. We compared the genotypes from the Miller et. al. [48] strain to our strain at ~93.7 million sites and found that 93.5 million sites (~99.8%) were homozygous reference in both datasets, while at 215,000 sites, both strains had the same heterozygous genotype. For another 37,000 sites, one strain was identified as homozygous and the other heterozygous. The strains were in complete disagreement (i.e. they were homozygous for different alleles) at only 3 sites. From this analysis, we concluded that the *D. triauraria* strain mislabeled *D. kikkawai* was in fact the same strain we sequenced.

### 2.2 Genome Synteny

We next sought to identify synteny blocks and assess the degree of chromosomal rearrangements between *D. melanogaster* and *D. triauraria*. We created an orthology map between the genome assemblies for these two species using *Mercator* [11] and identified a total of 991 synteny blocks with average size of ~117 Kb in *D. melanogaster* and ~140 Kb in *D. triauraria*. The larger size of the synteny blocks in *D. triauraria* is consistent with the larger genome size for this species. We visualized synteny by using the promer tool from the *MUMmer* pipeline [36] to produce a dotplot (Figure 2c), which shows that there have been extensive chromosomal rearrangements since the divergence of these two species, with the majority of rearrangements occurring within Muller elements.

### 2.3 TAD Boundary and Domain Annotation

In order to determine how the large number of chromosomal rearrangements present between these two species has affected 3D genome organization, we identified TAD boundaries as well as complete contact domains in both species. We used *HiCExplorer* [55], which was developed using Drosophila Hi-C data, to identify TAD boundaries at 5 kb resolution for both *D. melanogaster* and *D. triauraria*. The total number of Hi-C read pairs for each dataset are reported in Table S2. *HiCExplorer* calculates the TAD separation score for each bin in the genome and identifies TAD boundaries as those bins whose score shows significantly larger contact insulation compared to neighboring bins. We used a bin size of 5 kb and found that the TAD separation scores were highly correlated between replicate datasets for each species (Spearman’s rho: 0.995 [*D. melanogaster*] and 0.990 [*D. triauraria*] (Figure S2a,b). We also found that the majority of predicted boundaries were identified in each replicate independently (74% [*D. melanogaster*] 70% [*D. triauraria*]). We refer to the boundaries that were identified in both replicates as high confidence boundaries, and those identified in only one of the two replicates as low confidence boundaries. In total, we identified 701 and 843 high confidence TAD boundaries for *D. melanogaster* and *D. triauraria*, respectively, and 249 and 355 low confidence boundaries (Table S3).

*HiCExplorer* [55] links TAD intra-boundary regions together into full contact domains. Similar to our approach with boundary elements, we identified contact domains that were found independently in both replicate datasets as high confidence domains and those found only in one replicate as low confidence domains. In total, we identified 552 and 639 high confidence TAD domains for *D. melanogaster* and *D. triauraria*, respectively, and 593 and 811 low confidence domains (Table S3).

### 2.4 Boundary Motif Enrichment

In Drosophila, TAD boundaries are highly enriched for motifs recognized by the insulator proteins M1BP and BEAF-32 [55]. To validate boundary calls made by *HiCExplorer* [55], we used *Homer* [27] software to search the identified boundaries for enriched sequence motifs. Boundaries from both species were enriched for motifs recognized by M1BP (p = 1*e* − 17, [*D. melanogaster*], p = 1*e* − 42 [*D. triauraria*]) and BEAF-32 (p = 1*e* − 18 [*D. melanogaster*], p = 1*e* − 15 [*D. triauraria*]), which supports the accuracy of our boundary calls (Figure S3).

### 2.5 Domain and Boundary Conservation

We assessed the evolutionary conservation of TAD boundaries between *D. melanogaster* and *D. triauraria* by lifting over the high confidence *D. melanogaster* boundary coordinates to the *D. triauraria* genome coordinates. We created a whole-genome alignment between the two genome assemblies using *Cactus* [3] and performed the coordinate liftovers using the *halLiftover* [28] utility. We considered boundaries to be orthologous when high confidence boundary regions lifted over from *D. melanogaster* to *D. triauraria* overlapped either a high or low confidence boundary that was independently identified in *D. triauraria*. Out of a total of 701 boundaries identified in *D. melanogaster*, 654 were successfully lifted over to a corresponding region in *D. triauraria*. Of the lifted over boundaries, 492 (~75%) are orthologous between the two species and 163 (~25%) are melanogaster-specific (Table 1, Figure S4a). Our results suggest that the sequences that specify TAD boundaries are highly conserved over evolutionary time.

**Table 1:** Summary of results from *D. melanogaster* to *D. triauraria* liftover analysis. Percentages of “Unique lifted over to *D. triauraria*” represent number out of total boundaries or domains. Percentages of orthologous and non-orthologous boundaries represent number out of “Unique lifted over to *D. triauraria*”.

| Category | Boundaries | Domains |
| --- | --- | --- |
| Total in *D. melanogaster* | 701 | 552 |
| Unique lifted over to *D. triauraria* | 654 (93%) | 544 (99%) |
| Orthologous | 492 (75%) | 134 (25%) |
| Non-orthologous | 163 (25%) | 410 (75%) |

There are two possibilities that would explain the high degree of boundary conservation in spite of the large number of chromosomal rearrangements between these species. One possibility is that boundaries are conserved because chromosomal rearrangements occur in such a way that TADs are shuffled as intact units (see Model 1, Figure 1). The other possibility is that the sequences that specify boundaries remain conserved while chromosomal rearrangements shuffle these sequence elements in ways that lead to widespread TAD reorganization (see Model 2, Figure 1). To differentiate between these possibilities, we identified orthologous contact domains between these two species. Similar to our approach with boundaries, we considered contact domains to be orthologous when high confidence domain regions from *D. melanogaster* lifted over as a continuous block (allowing for internal rearrangements) to *D. triauraria* and overlapped either a high or low confidence TAD domain that was independently identified in *D. triauraria*. We required that the domains were reciprocally overlapping by at least 90% of their lengths.

Out of a total of 552 domains identified in *D. melanogaster*, 544 were successfully lifted over to a corresponding region in *D. triauraria*. Of the lifted over domains, we found that 134 (25%) are orthologous between the two species, whereas 410 (75%) of the *D. melanogaster* TADs do not show a one-to-one relationship with a *D. triauraria* TAD (Table 1, Figure S4a). Most of the non-orthologous TADs are due to cases where TADs have been split (i.e. a contiguous *D. melanogaster* domain lifts over to multiple, discontiguous regions in *D. triauraria*) (Figure 3b). Of the orthologous domains, 84 (~63%) also shared orthologous boundary regions.

**Figure 3:**
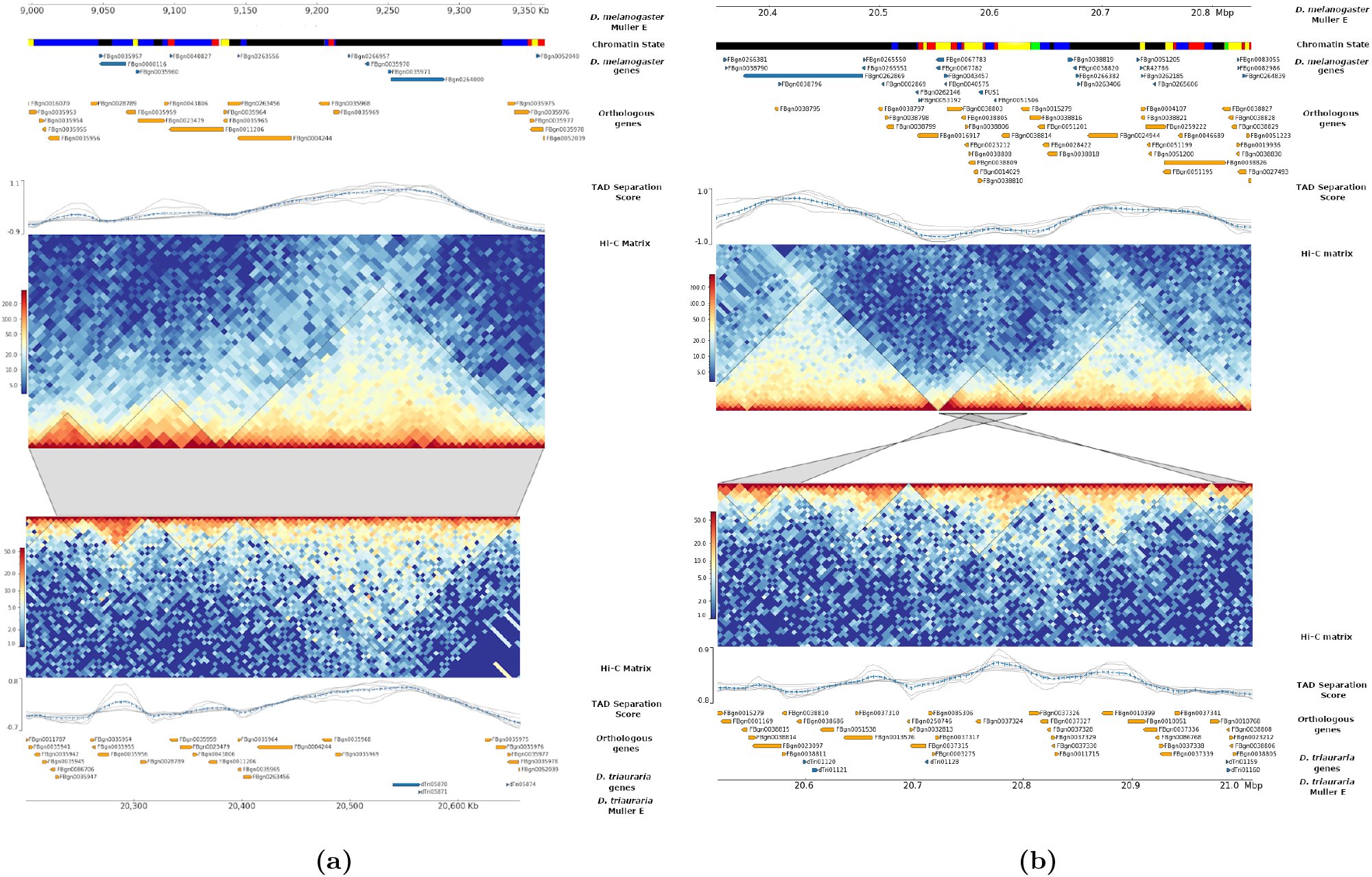
Visualization of orthologous and non-orthologous TADs. a) Three orthologous, conserved TADs on Muller E between *D. melanogaster* (upper heatmap) and *D. triauraria* (lower heatmap). b) Non-orthologous, split TAD on Muller E between *D. melanogaster* (upper heatmap) and *D. triauraria* (lower heatmap). The black triangles on the contact matrices show the locations of TADs. The chromatin state annotations are based on Filion et. al. [20]. The chromatin classifications are as follows: *black*: inactive, *blue*: Polycomb-repressed heterochromatin, *green*: constitutive heterochromatin, *red*: dynamic active, *yellow*: constitutive active. Orthologous genes are labeled by their FlyBase IDs and Hi-C matrices were generated by *HiCExplorer*. The grey blocks connecting matrices indicate syntenic regions. In b) gene tracks show that genes such as FBgb0038805, FBgn0038806, and FBgn0038814 are split between different TADs in *D. triauraria* and are in the reverse orientation compared to *D. melanogaster*.

The Drosophila X chromosome has previously been shown to accumulate chromosomal rearrangements at a faster rate compared to the autosomes [4]. We found a similar pattern for *D. melanogaster* and *D. triauraria*, where the median size of synteny blocks is significantly lower for the X chromosome compared to the autosomes (Figure S5, Wilcoxon test p = 1.35*e*−12). We also found that the proportion of orthologous TADs on the X chromosome is reduced relative to the autosomes (Figure S5, Fisher’s Exact Test p = 0.014), consistent with increased structural divergence of the X chromosome leading to increased TAD reorganization.

Given that only 25% of domains are orthologous between the two species, we conclude that chromosomal rearrangements have reorganized the majority of TADs present in each of these species since their common ancestor. Comparing the boundary and domain data, the rate of conservation of TAD boundaries is much higher than domains (Fisher’s Exact Test p = 4.78*e* − 63) (Figure S4). Our results suggest that Model 2 (Figure 1) is the most likely scenario for TAD evolution: boundary regions are conserved but contact domains are reorganized between species. For consistency, we repeated these analyses by performing the liftover in the opposite direction, from *D. triauraria* to *D. melanogaster*, and obtained similar results (Table S4, Figure S4b).

### 2.6 Gene Expression

Enhancer-promoter contacts regulate gene expression. We hypothesized that TADs rearranged in *D. triauraria* compared to *D. melanogaster* might reorganize enhancer-promoter contacts and result in altered gene expression profiles. We performed RNA-seq on replicate datasets for each species and used the *DESeq* R package [1] to identify differentially expressed genes between the two species. A total of 964 differentially expressed genes were identified (Figure S6). We then compared the expression of genes within orthologous and non-orthologous TAD domains between the two species and found that, while nonconserved TADs show a slightly higher percentage of differentially expressed genes (10.5% versus 9.1%), this difference is not significant (Figure 4, Fisher’s Exact Test p = 0.151). We did, however, find that genes in orthologous TADs are expressed at significantly lower levels than those in non-orthologous TADs (Wilcoxon test p = 6.7*e* − 05) (Figure 4a). We also found that orthologous TADs are enriched for homeobox domain-containing genes (FlyMine protein domain enrichment test, Benjamini-Hochberg corrected p = 0.02) [45] and, in comparison to the genes within non-orthologous TADs, are also highly enriched for genes predicted to be regulated by Polycomb-group proteins (Fisher’s Exact Test p = 2.6*e* − 5) [7].

**Figure 4:**
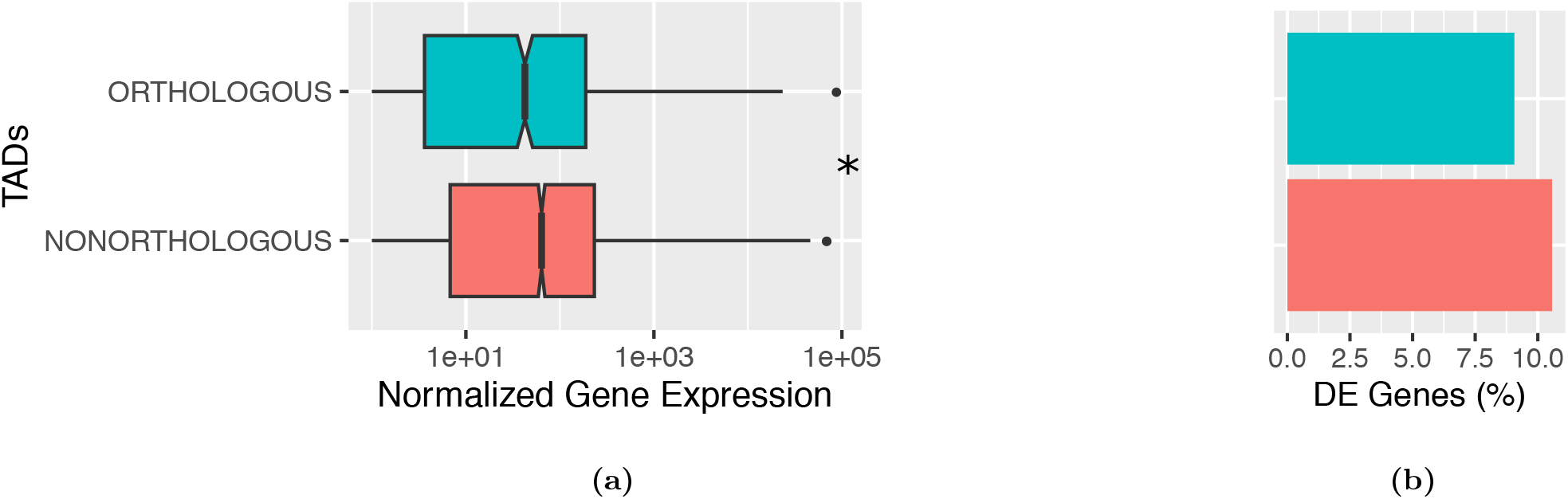
Gene expression in orthologous versus non-orthologous TADs. a) Genes within orthologous TADs are expressed at significantly reduced levels compared to non-orthologous TADs, consistent with polycomb-repression (Wilcoxon test p = 6.7*e* − 05) b) Orthologous TADs contain slightly fewer differentially expressed genes compared to non-orthologous TADs (9.1% versus 10.5%), however this difference is not statistically significant (Fisher’s Exact Test p = 0.151). Differentially expressed genes were identified using the *DESeq2* R software package [1].

### 2.7 Chromatin State

Our finding that orthologous TADs are enriched for Polycomb-repressed genes, which are characterized by the *blue* chromatin state described by Filion et al. [20], led us to compare chromatin states between genes in orthologous and non-orthologous TADs. We quantified the number of genes in each of five chromatin states within orthologous and non-orthologous TAD regions (Table S5). Orthologous TADs show significant enrichment of the *black* (transcriptionally silent) and *blue* (Polycomb-repressed) chromatin states and significant depletion of the *green* (constitutive heterochromatin) and *yellow* (constitutively active) chromatin states, compared to non-orthologous TADs (Fisher’s Exact Test ps: 1.225*e* − 25 [*black*], 4.322*e* − 4 [*blue*], 1.552*e* − 15 [*green*], 8.375*e* − 23 [*yellow*]) (Figure 5). The enrichment of silenced and repressed chromatin states are in line with our result revealing that gene expression is significantly lower in orthologous TADs (Figure 4a). The chromatin state tracks in Figure 3a and 3b also support our findings. The majority of genes in the conserved TADs in Figure 3a are black and blue, while the genes within the split TAD in Figure 3b are predominantly *red* and *yellow*. These results largely mirror the chromatin states of the ancient and highly conserved contact domains identified by Harmston et. al. [26], which contain clusters of conserved non-coding elements and developmental genes.

**Figure 5:**
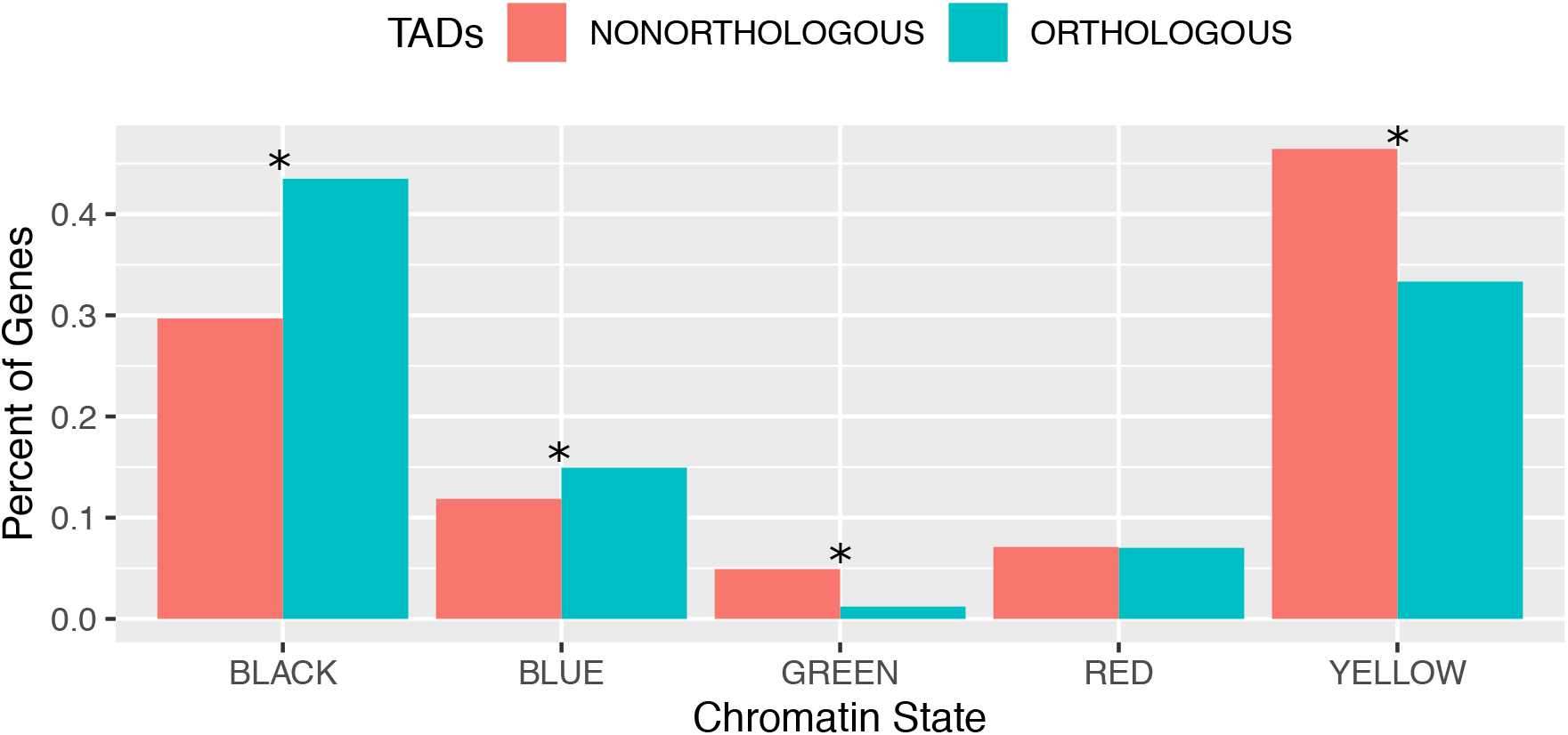
Orthologous TADs are enriched for transcriptionally silent and polycomb-repressed genes. The bar plot shows the percent of genes in each chromatin state (defined by [20]) in orthologous versus non-orthologous TADs. Orthologous TADs are significantly enriched for the black (inactive) and blue (Polycomb-repressed) chromatin states and significantly depleted of the green and yellow chromatin states. Asterices *ú* indicate significant differences as calculated by Fisher’s Exact Test (p-values: Black = 1.225*e* − 25; Blue = 4.322*e* − 4; Green = 1.552*e* − 15; Yellow = 8.375*e* − 23; Red = 0.398.)

## 3 Discussion

In this study, we sought to examine the evolutionary conservation of 3D genome organization in Drosophila. We selected *D. melanogaster* and *D. triauraria* for this comparison because they are separated by ~15 million years of evolution [50]. We predicted that this level of divergence would be long enough that large-scale chromosomal rearrangements would have occurred between the two species but short enough that conservation at the nucleotide level would allow for an accurate whole-genome alignment. We used a combination of nanopore and Illumina Hi-C sequencing data to improve a recently published *D. triauraria* genome assembly produced from relatively low-coverage (depth 18.8x) nanopore sequencing data [48]. We have previously shown that Hi-C data can be used to scaffold Drosophila nanopore contigs with high accuracy, and even correct contig misassemblies [16]. We used our Hi-C data to scaffold the *D. triauraria* nanopore contigs and our improved *D. triauraria* assembly resulted in chromosome-length scaffolds highly enriched for genes corresponding to a single Muller element (Figure S1), further supporting the efficacy of this approach. We were able to align ~87% of our *D. triauraria* assembly to the *D. melanogaster* reference assembly and we found extensive chromosomal rearrangements (Figure 2c), consistent with our initial prediction that *D. triauraria* and *D. melanogaster* represent an ideal species pair for use in a comparative study of 3D genome organization.

Previous research has yielded conflicting results regarding the evolutionary conservation of TAD domains. In theory, TADs should be under strong purifying selection due to their role in preventing aberrant gene-enhancer interactions. Therefore, we expected that entire TAD contact domains, including boundary regions, would be conserved (i.e. Model 1, Figure 1). However, we found that TADs themselves diverge much faster than their boundary elements, which suggests that conserved boundary elements are shuffled by chromosomal rearrangements, resulting in reorganization of TAD architecture (see Model 2, Figure 1). This finding is similar to several recent studies suggesting that TADs may actually diverge relatively rapidly and that TAD reorganization is not necessarily associated with divergence in gene expression [19, 24, 39, 47, 10].

Why, then, have other studies concluded that 3D genome organization is highly conserved? One possibility is that differences in the measurements used to assess conservation will produce contrasting results. For example, some studies have based their conclusion of strong conservation on a statistical correlation of contact frequencies between species, without identifying TADs [68] or on a statistical association between synteny breakpoints and TAD boundaries [57, 38]. Renschler et. al. [57] report that 3D genome architecture is conserved across the Drosophila species *D. melanogaster*, *D. virilis*, and *D. busckii* based on a significant association between synteny breakpoints and TAD boundaries, yet find that only 10% of identified TADs were conserved across all three species. An accurate assessment of 3D genome conservation should identify orthologous TADs while also comparing the rate of TAD divergence to other regulatory phenotypes [19]. Eres et. al. [19] report that human-chimp TADs diverge much more rapidly than any than any other regulatory phenotype. Similarly, we find that Drosophila TADs diverge much more rapidly than both boundary elements and gene expression (75% divergent TADs versus 25% divergent boundaries and 11% DE genes).

It is also possible that inconsistent assessments of TAD conservation are due to differences in the approaches used to identify TADs in the first place. Previous studies have identified inconsistencies in TAD-calling software packages [71] and have raised the possibility that TAD conservation results may depend on the direction of the liftover comparison [19]. For example, some studies report conservation estimates by first calling TADs in the species for which they have more data and then identifying the orthologous domains in the species for which they have less data [19, 56, 12, 57]. When reversing the analysis the conservation rate can be reduced by up to 25% [19]. However, in our study, we used biological replicates to demonstrate that the identification of TAD boundaries and TAD units is highly reproducible. We also performed our analysis of TAD conservation in both directions (i.e. from *D. melanogaster* to *D. triauraria* and vice versa) and obtained similar results regardless of the direction of comparison (Figure S4). Furthermore, our estimates of conservation, if biased at all, should be biased towards inferring higher levels of conservation. We only considered TADs for our liftover step if they were independently identified in both biological replicates, which should enrich for stronger TADs. Furthermore, after liftover, we considered the TAD to be orthologous if it overlapped *either* a strong (i.e. high-confidence) TAD *or* a weak TAD (i.e. low-confidence TAD identified in only a single replicate). We also did not require orthologous TADs to have orthologous boundaries. Instead, they were only required to have a reciprocal overlap of at least 90% of their lengths. We would expect these relatively low-stringency criteria to potentially result in an over-estimate of TAD conservation, yet we still only find ~25% of TADs to be orthologous between species.

One potential function of TADs is to reduce aberrant enhancer-promoter contacts by constraining 3D interactions to specific segments of the chromosome. Some genome editing experiments have shown that deletion of TAD boundaries results in the fusion of neighboring TADs and is accompanied by misregulation of gene expression [49, 43]. However, other studies show that TAD disruption via boundary removal or chromosomal rearrangement does not affect gene regulation [10, 24]. We compared the gene expression profiles within orthologous TADs to non-orthologous TADs. Slightly (~2%) more of the genes in non-orthologous domains are differentially expressed, however this difference is not significant (Fishers Exact Test p = 0.151). It is possible that the lack of association between TAD reorganization and gene expression divergence is due to compensatory mutations that have occurred since the common ancestor of these two species. The relatively large divergence time between these two species could have allowed for the fixation of mutations that compensate for suboptimal gene expression patterns caused by TAD reorganization. Alternatively, it is also possible that, in the relatively compact Drosophila genome, most gene regulatory sequences are directly adjacent to the genes they regulate and are therefore rarely separated by chromosomal rearrangements or TAD reorganization [39].

Our chromatin evaluation revealed that orthologous TADs are significantly enriched for inactive and Polycomb-repressed chromatin states, as well as homeobox-domain containing genes and genes predicted to be regulated by Polycomb-group (PcG) proteins. PcG proteins play an important role in regulation of developmental genes [7], bringing our results in line with previous studies showing that TADs enriched for developmental genes are more likely to be conserved [26]. Developmental genes are often surrounded by clusters of conserved noncoding elements (CNEs) [59]. These clusters of CNEs have been termed genomic regulatory blocks (GRBs) [17] and are highly-conserved [59, 17]. It has been recently discovered that GRBs coincide strongly with a subset of highly-conserved TADs in vertebrates and invertebrates ranging from Drosophila to humans [26, 35]. Most enhancers in Drosophila are within 10 kb of the promoter they regulate [9], whereas developmental genes have more complex regulatory landscapes which can extend to hundreds of kilobases [9].

Our results show that TAD structures diverge rapidly and this divergence in 3D organization is not associated with gene expression divergence. Instead, highly conserved TADs are enriched for Polycomb-repressed genes, lending more support for an important link between Polycomb-repressed chromatin and 3D genome organization in Drosophila [18]. One possibility that would explain the apparent contradiction between studies that have shown that gene misregulation is associated with the engineered disruption of a specific TAD versus genome-wide studies that have found little to no association between TAD reorganization and differential gene expression, is that there are different subtypes of TADs with variable tolerances for disruption. Disruption of some types of TADs, such as those containing developmental genes surrounded by clusters of conserved non-coding regulatory elements, may detrimentally affect gene expression profiles and are therefore highly conserved. Other types of TADs, such as those containing compact genes with adjacent regulatory elements, could potentially be altered without any major effects on gene expression, since chromosomal rearrangements will be more likely to shuffle the gene and its regulatory sequences together as single unit. These TADs may be more likely to diverge quickly between species. If this is true, previous studies reporting contradictory effects of TAD rearrangement on gene expression may simply be due to the differences in the subtypes of TAD being tested. Future work involving experimental disruption of an unbiased sample of TADs would allow for testing of this prediction.

## 4 Methods

### 4.1 *D. triauraria* Genome Sequencing

Using the Qiagen DNAeasy Blood and Tissue Kit, we extracted DNA from ~30 *D. triauraria* adult females strain 14028-0651.00 (National Drosophila Species Stock Center at Cornell). We used the Oxford Nanopore Technologies (ONT) SQK-LSK 108 library preparation kit to construct three PCR-free libraries according to the ONT 1D Genomic DNA by Ligation protocol. Each library was sequenced on a MinION r9.4 flow cell. Raw signal data were basecalled using the ONT *Albacore* software package with default parameters.

### 4.2 Hi-C Chromosome Conformation Capture

*D. triauraria* and *D. melanogaster* strains were maintained in population cages on molasses agar with yeast paste. Embryos (8-16 h) for each species were collected and dechorionated in 50% commercial bleach for 2.5 min. Nuclei were isolated from ~1 g of embryos and fixed in 1.8% formaldehyde for 15 minutes according to the protocol in Sandmann et. al. [60]. Two replicate Hi-C libraries were constructed for each species using the *in situ* DNase Hi-C protocol described by Ramani et. al. [54]. Libraries were sequenced on an Illumina HiSeq 2500 machine.

### 4.3 RNA-seq

Approximately 0.02 g fresh embryos were collected using the same approach as for Hi-C libraries and homogenized in 300 *μL* 1x DNA/RNA Shield. Fifty pairs of testes were dissected from 3-5-day old mated males and 10 pairs of ovaries from 3-5-day old mated females. Dissections were performed in 1X PBS and then immediately transferred into 200 *μL* RNAlater solution. Two hundred *μL* of ice-cold 1X PBS was added to each sample and they were centrifuged at 5000g for 1 min at 4°C. After removing the supernatant, 300 *μL* 1× DNA/RNA shield was added and samples were homogenized immediately using an electric pestle.

RNA was extracted using the Quick-RNA Plus Kit (R1057) from Zymo Research. Samples were incubated at 55°C with 30 *μL* PK Digestion Buffer and 15 *μL* Proteinase K for at least 30 minutes. Column-based size selection was used to obtain >200 nt purified total RNA. MRNA-seq libraries were constructed from the total RNA using the NEBNext Poly(A) mRNA Magnetic Isolation Module (E7049) and the NEBNext Ultra II Directional RNA Library Prep Kit for Illumina (E7760) using 1 *μg* total RNA with fragmentation for 7 min at 94°C and first strand cDNA synthesis via incubation for 45 minutes at 42°C. Library quality was assessed on a Bioanalyzer. For the embryo samples, two biological replicate libraries were prepared for *D. triauraria* strain 14028-0651.00 (National Drosophila Species Stock Center at Cornell) while *D. melanogaster* strains RAL-379 and RAL-732 from the Drosophila Genetic Reference Panel (DGRP) [46] were used as biological replicates for *D. melanogaster*.

### 4.4 *D. triauraria* Genome Assembly and Annotation

The basecalled *D. triauraria* nanopore reads were combined with the nanopore reads from Miller et. al. [48] and assembled using *Canu* [33]. *Purge Haplotigs* [58] was used to identify and collapse heterozygous contigs where each haplotype was assembled separately. The assembly was then polished using the raw nanopore signal data with *Nanopolish* [42]. Finally, uninformative Hi-C Illumina sequences (i.e. those that do not contain a ligation junction) were used as single-end reads to further polish the assembly with *Pilon* [69]. To confirm that our data were from the same *D. triauraria* strain as the data generated by Miller et. al. [48], we aligned our uninformative *D. triauraria* Hi-C reads and the *D. triauraria* Illumina data from Miller et. al. (SRA project PRJNA473618) [48] as single-end reads using *bowtie2* (version 2.2.9) [37] with default parameters. We then called SNPs using *Freebayes* (version 1.2.0) [23] with the parameters –*no-indels –no-mnps –no-complex −0 –report-monomorphic –use-best-n-alleles 4*.

The *Juicer* [15] and 3D-DNA [13] pipelines were used to scaffold the *D. triauraria* nanopore reads with Hi-C sequencing data. The *Juicebox* software package [14] was used to visualize contact matrices, assign chromosome boundaries, and export a finalized reference sequence for downstream analysis.

To assign the chromosome-length *D. triauraria* scaffolds to their corresponding Muller element (i.e. Muller A-F), we performed a translated BLAST search of our scaffolds using FlyBase r6.21 [66] *D. melanogaster* peptides as queries (Figure 2a,b).

To pre-process the RNA-seq data for *MAKER* [8], we first aligned the *D. triauraria* RNA-seq Illumina reads to the newly assembled *D. triauraria* reference genome using *HISAT2* [32]. Second, the *HISAT2* alignments were used to assemble mRNA transcripts with *stringtie* [51]. The *stringtie* transcripts were provided to *MAKER* along with *D. melanogaster* r6.26 peptides from FlyBase [66]. The *MAKER* control file is available via GitHub.

### 4.5 Genome Synteny

We identified synteny blocks between *D. melanogaster* and *D. triauraria* using *Mercator* [11] software. We visualized synteny using the promer tools from the *MUMmer* [36] pipeline to produce a dotplot comparison of the *D. melanogaster* and *D. triauraria* genomes.

### 4.6 Identifying TAD Boundaries and Domains

We removed adapter sequences from Hi-C reads for each species using *Trimmomatic* [5] and used a custom perl script to split reads that contain a ligation junction. We used BWA software [40] to align the split forward and reverse Hi-C reads to each species’ reference assembly (the *D.triauraria* assembly generated in this study and the *D. melanogaster* release 6 assembly from Flybase [66]). We used *HiCExplorer* version 2.2 [55] to create a normalized contact frequency matrix. To find TAD boundaries and domains for each species we ran the *hicFindTads* utility separately for each biological replicate. We used *Bedtools* [53] to identify overlapping boundaries and domains between replicates. Boundaries were required to overlap by at least one base pair in both replicates. Domains were required to have start and end coordinates within 5000 bp in both replicates. Boundaries and domains identified in both replicates were considered high confidence while those identified in one replicate are low confidence. We used a custom python script to calculate correlation coefficients between replicates for the TAD separation scores.

### 4.7 Defining and identifying orthologous TAD boundaries between *D. melanogaster* and *D. triauraria*

We softmasked the *D. melanogaster* and *D. triauraria* genomes using *Repeatmasker* [62] and aligned them using *Cactus* [3] to generate a hal file. We input the high confidence boundaries for *D. melanogaster* to *halLiftover* [28] to identify the corresponding genomic coordinates in *D. triauraria*. *HalLiftover* reports contiguous liftover coordinates as separate features if they include short indels. We therefore merged ‘lifted-over’ boundary locations that were within 5000 bp of each other into a single feature. Lifted over boundaries less than 500 bp in size were excluded from further analysis. We considered lifted-over boundaries from *D. triauraria* that were located less than 5 kb from either a high or low confidence boundary in *D. melanogaster* to be orthologous boundaries. Lifted-over boundaries from *D. triauraria* that were not identified as boundaries in *D. melanogaster* were considered non-orthologous. We implemented the same pipeline for the reverse comparison, from *D. triauraria* to *D. melanogaster*.

### 4.8 Boundary Motif Enrichment

We used *Homer* [27] software to search for enriched motifs with the high-confidence boundary sequences for both *D. triauraria* and *D. melanogaster*. We split each genome assembly into 5 kb sequences for use as the background dataset.

### 4.9 Defining and identifying orthologous domains between *D. melanogaster* and *D. triauraria*

To assess domain conservation between *D. melanogaster* and *D. triauraria* we used *halLiftover* [28]. *HalLiftover* will report lifted-over coordinates as separate features if there are species-specific indels, transposon insertions, or chromosomal rearrangements. In order to combine contiguously lifted-over segments that were separated by species-specific indels, TE insertions, or intra-TAD rearrangements, we merged lifted-over features separated by less than 20 kb. Lifted over features less than 5000 bp were excluded from further analysis. After merging, the lifted-over domains in *D. triauraria* that reciprocally overlapped a *D. melanogaster* high or low confidence domain (>90%) were considered orthologous domains. Lifted-over domains in *D. triauraria* that did not meet the reciprocal overlap criteria with a *D. melanogaster* domain were considered non-orthologous. To identify orthologous domains between the two species that also share boundaries, we required the *D. triauraria* lifted-over endpoints to lie within 5 kb of the *D. melanogaster* orthologous TAD boundaries. We implemented the same pipeline for the reverse comparison, from *D. triauraria* to *D. melanogaster*.

### 4.10 Gene Expression

Stranded embryo RNA-seq data were aligned to their respective genomes using *hisat2* (version 2.1.0) [32] with parameters *–dta –max-intronlen 50000 –rna-strandness RF*. Per-gene raw read counts were generated using *htseq-count* (version 0.11.2) [2] with parameters -*i Parent -f bam -r pos -s reverse -a 20 –nonunique none*. Our *MAKER* [8] gene models were used for *D. triauraria* and the FlyBase r6.21 [66] gene models were used for *D. melanogaster*. One-to-one gene orthologs were identified using our *Mercator* [11] orthology map and differentially expressed genes were identified using the *DESeq2* R software package [1].

### 4.11 Chromatin State

We used the chromatin state annotations from Filion et. al. [20] to assign each *D. melanogaster* gene to one of five chromatin states (*black*, *blue*, *green*, *red*, *yellow*). Genes were assigned to chromatin states based on the state that covered the largest proportion of the gene (including introns) and we counted the number of genes from orthologous versus non-orthologous TADs that for each of the five chromatin states.

### 4.12 Data Availability

Nanopore and Illumina data generated for this project are available at the National Center for Biotechnology Information (https://www.ncbi.nlm.nih.gov/) under BioProject PRJNA627893. Complete analysis pipelines and all custom scripts described in this project can be found on GitHub at https://github.com/Ellison-Lab/TADs.git.

## 5 Acknowledgements

The authors acknowledge the Office of Advanced Research Computing (OARC) at Rutgers, The State University of New Jersey for providing access to the Amarel cluster and associated research computing resources that have contributed to the results reported here. Stocks obtained from the Bloomington Drosophila Stock Center (NIH P40OD018537) and the National Drosophila Species Stock Center were used in this study.

