## Supplementary Tables and Figures for "3D genome evolution and reorganization in the *Drosophila melanogaster* species group"

#### Tables

| | Nanopore Sequencing (this study) | Nanopore Sequencing (combined) | 200M Genome Assembly | Post Quality Control and Hi-C Scaffolding |
|---|---|---|---|---|
| Number of Sequences/Contigs | 633,844 | 1,043,600 | 1,098 | 302 |
| Sum of Sequence/Contig Length | 6,514,819,227 | 10,462,257,773 | 268,538,400 | 198,798,533 |
| Average Sequence/Contig Length | 10,278 | 10,025 | 244,570.49 | 658,273 |
| Median Sequence Length | 7,405 | 6,549 | 60,582.50 | 24,380 |
| Longest Sequence Length | 123,032 | 238,837 | 7,592,683 | 37,891,417 |
| N50 | – | – | 1,274,151 | 34,538,105 |

**Table S1: Combined *D. triauraria* sequencing and assembly data.** This study and Miller et al. [48])

| Species | Replicate | Total number of read pairs | Pairs mappable, unique and high quality | Pairs used |
|---|---|---|---|---|
| *D. melanogaster* | 1 | 126758371 | 65470274 | 41063921 |
| *D. melanogaster* | 2 | 157512176 | 71652238 | 45294778 |
| *D. triauraria* | 1 | 56128496 | 18491258 | 14551364 |
| *D. triauraria* | 2 | 57584713 | 19513854 | 15023262 |

**Table S2: Total number of Hi-C read pairs for each species and replicate, the number of pairs considered and used by *HiCExplorer* [55].**

| Species | Boundaries | | Domains | |
|---|---|---|---|---|
| | High Confidence | Low Confidence | High Confidence | Low Confidence |
| *D. melanogaster* | 701 | 249 | 552 | 593 |
| *D. triauraria* | 843 | 355 | 639 | 811 |

**Table S3: Counts of high and low confidence TAD boundaries and domains identified in *D. melanogaster* and *D. triauraria*.**

| **Category** | **Boundaries** | **Domains** |
|---|---|---|
| Total in *D. triauraria* | 834 | 639 |
| Unique lifted over to *D. melanogaster* | 798 | 637 |
| Orthologous | 503 | 177 |
| Non-orthologous | 295 | 460 |

**Table S4: Summary of results from *D. melanogaster* to *D. triauraria* liftover analysis.**

| **Chromatin State** | **Orthologous TADs: number of genes** | **Non-orthologous TADs: number of genes** |
|---|---|---|
| Black | 719 | 2156 |
| Blue | 247 | 861 |
| Green | 20 | 357 |
| Red | 116 | 516 |
| Yellow | 551 | 3373 |

**Table S5: Number of genes of each chromatin state in orthologous and non-orthologous TADs.**

### Figures

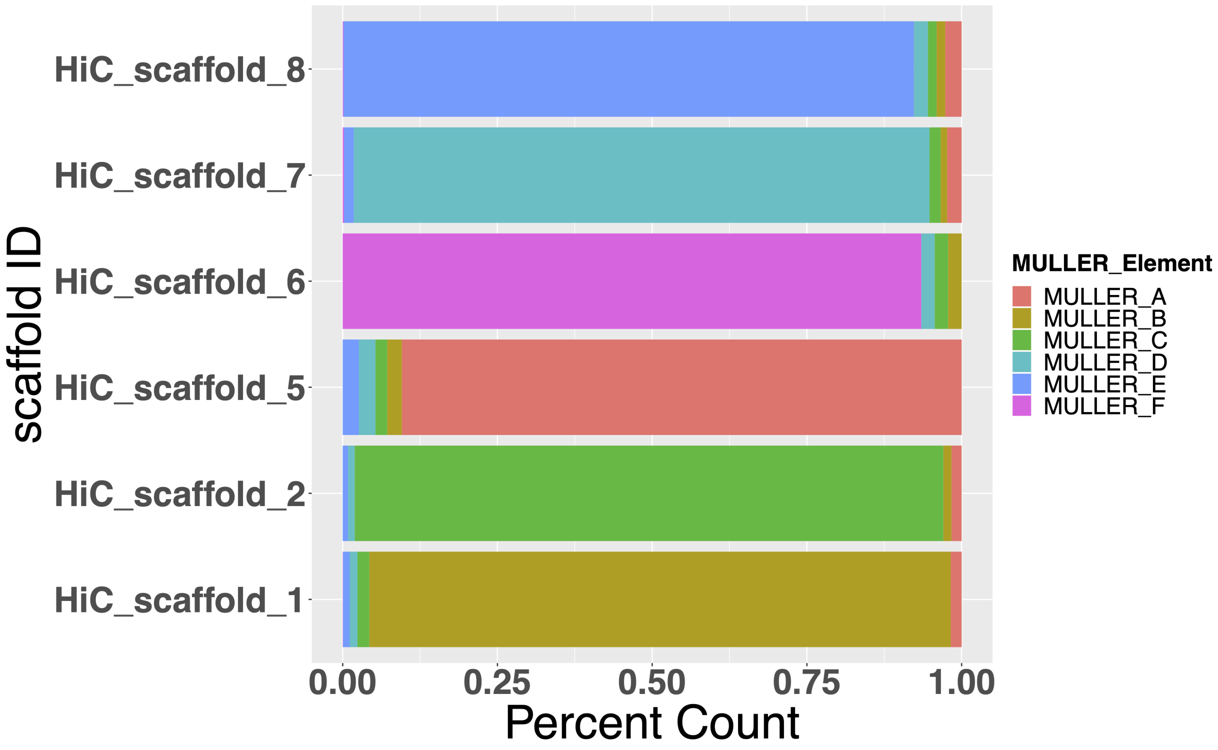


**Figure S1: Muller Element genes per *D. triauraria* megascaffold.** Percent of *D. melanogaster* genes corresponding to each of the *D. triauraria* chromosome-length scaffolds. Each scaffold is enriched for genes belonging to a single Muller element.

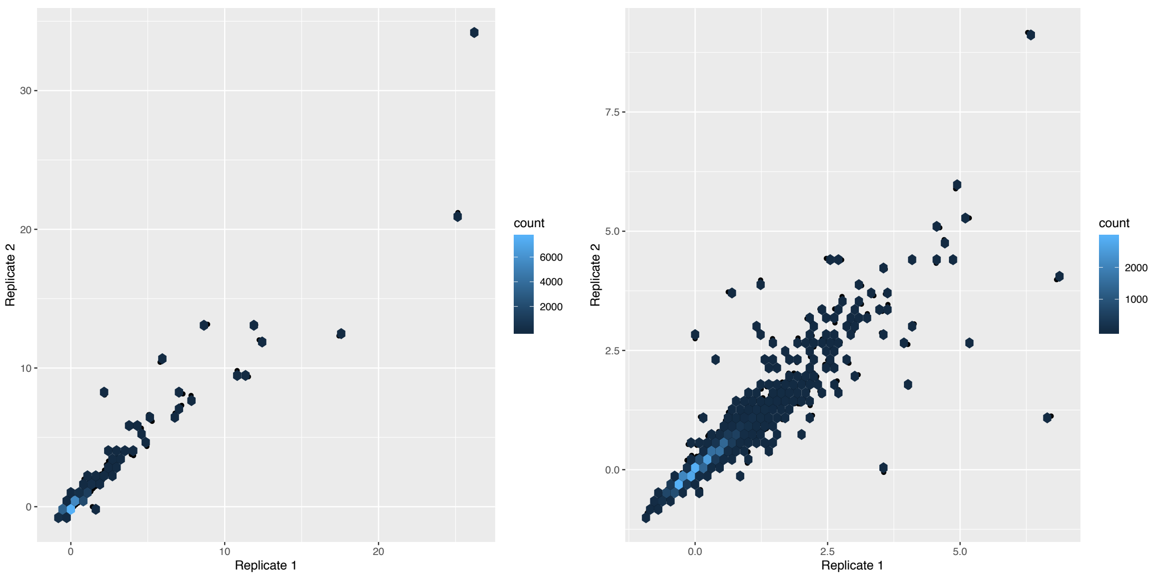


(a) (b)

**Figure S2: Scatterplots depicting correlation between TAD separation score between replicate datasets.** TAD separation scores from each replicate plotted for a) *D. melanogaster* (Spearman's rho: 0.995) and b) *D. triauraria* (Spearman's rho: 0.990).

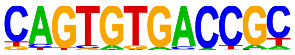

**(a)**

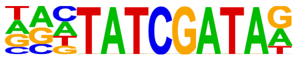

**(b)**

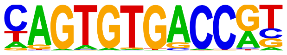

**(c)**

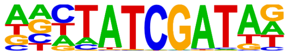

**(d)**

**Figure S3: Sequence logo diagrams for enriched motifs at TAD boundaries.** Motifs for a) *D. melanogaster* M1BP ($p = 1e-17$), b) *D. melanogaster* BEAF-32/DREF ($p = 1e-18$), c) *D. triauraria* M1BP ($p = 1e-42$), and d) *D. triauraria* BEAF-32/DREF ($p = 1e-15$). *Homer* [27] software found these sequence motifs to be enriched at TAD boundaries in *D. melanogster* and *D. triauraria*. BEAF-32 and DREF binding motifs are almost identical and both recognized by the BEAF-32 insulator protein [25].

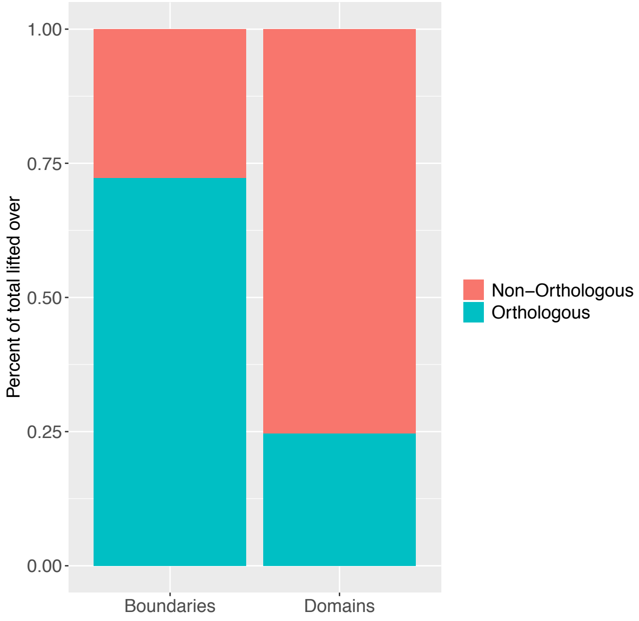


**(a)**

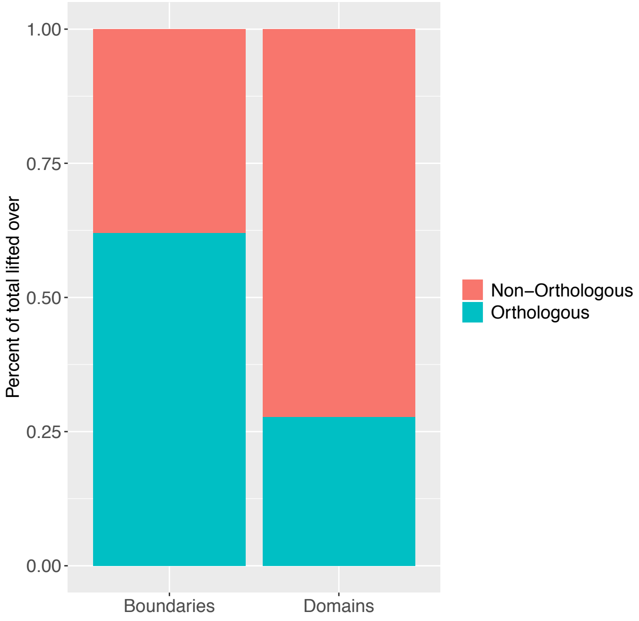


**(b)**

**Figure S4: Rate of boundary orthology is higher than domain orthology for both directional analyses.** Percent of unique lifted over TAD domains and boundaries that are orthologous and non-orthologous in a) *D. melanogaster* to *D. triauraria* liftover analysis, and b) *D. triauraria* to *D. melanogaster* liftover analysis.

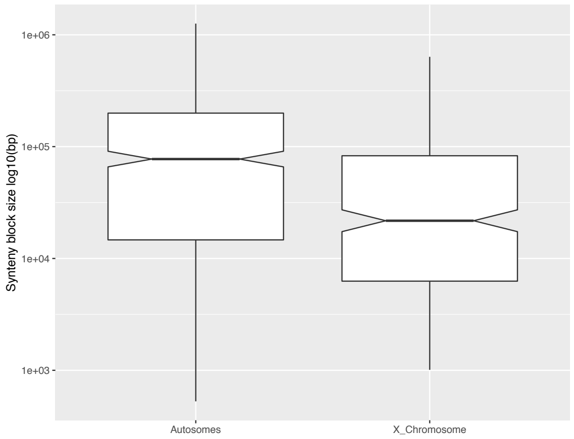


**(a)**

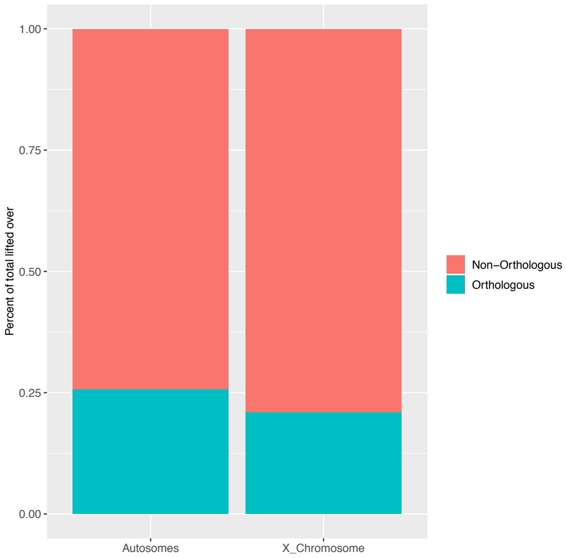


**(b)**

**Figure S5: The X chromosome has more chromosomal rearrangements and fewer orthologous TADs, compared to autosomes.** (a) The size of *D. triauraria* versus *D. melanogaster* synteny blocks is significantly reduced on the X chromosome (Muller A) relative to the autosomes, indicating that the X has accumulated more chromosomal rearrangements (Wilcoxon test $p = 6.7e-05$). (b) The proportion of orthologous versus non-orthologous TADs is also significantly reduced on the X chromosome (Fisher's Exact Test $p = 0.0135$)

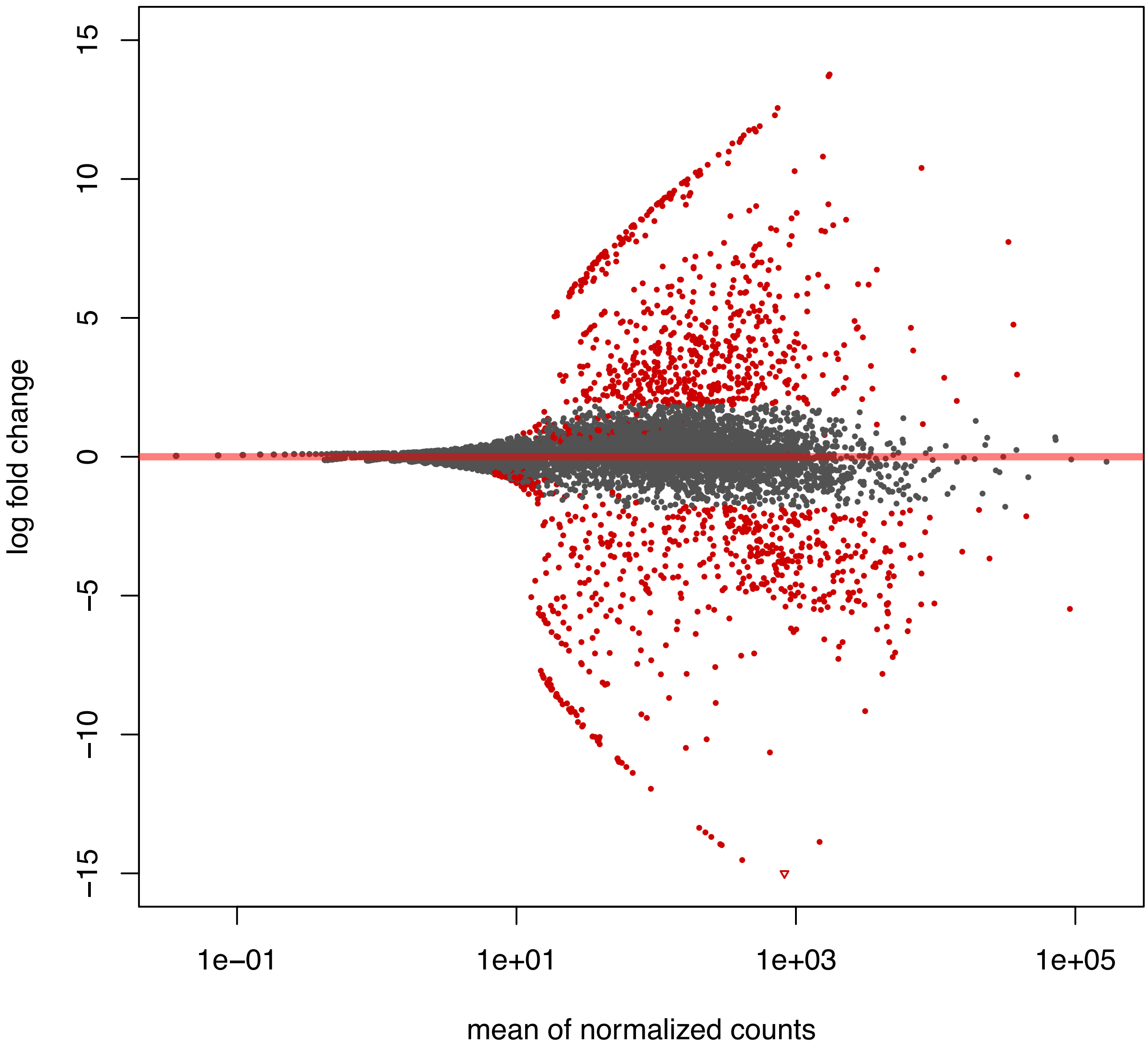


!h

**Figure S6: MA plot highlighting differentially expressed genes between *D. triauraria* and *D. melanogaster*.** Differentially expressed genes (adjusted p-value <= 0.05) between *D. melanogaster* and *D. triauraria* identified by *DEseq2* [1] are shown in red. Each point represents an orthologous gene pair between the two species. The plot was created using the *DEseq2* shrunken log2 fold changes which removes noise from low count genes.
